# KEEPING IN TOUCH WITH OUR HIDDEN SIDE

**DOI:** 10.1101/2021.11.23.469678

**Authors:** Benjamin Mathieu, Antonin Abillama, Malvina Martinez, Laurence Mouchnino, Jean Blouin

**Author notes:** Co-first authorship: Benjamin Mathieu and Antonin Abillama contributed equally to this work. Correspondance : Benjamin Mathieu.

## Abstract

Previous studies have shown that the sensory modality used to identify the position of proprioceptive targets hidden from sight, but frequently viewed, influences the type of the body representation employed for reaching them with the finger. The question then arises as to whether this observation also applies to proprioceptive targets which are hidden from sight, and rarely, if ever, viewed. We used an established technique for pinpointing the type of body representation used for the spatial encoding of targets which consisted of assessing the effect of peripheral gaze fixation on the pointing accuracy. More precisely, an exteroceptive, visually dependent, body representation is thought to be used if gaze deviation induces a deviation of the pointing movement. Three light-emitting diodes (LEDs) were positioned at the participants’ eye level at −25 deg, 0 deg and +25 deg with respect to the cyclopean eye. Without moving the head, the participant fixated the lit LED before the experimenter indicated one of the three target head positions: topmost point of the head (vertex) and two other points located at the front and back of the head. These targets were either verbal-cued or tactile-cued. The goal of the subjects (n=27) was to reach the target with their index finger. We analysed the accuracy of the movements directed to the topmost point of the head, which is a well-defined, yet out of view anatomical point. Based on the possibility of the brain to create visual representations of the body areas that remain out of view, we hypothesized that the position of the vertex is encoded using an exteroceptive body representation, both when verbally or tactile-cued. Results revealed that the pointing errors were biased in the opposite direction of gaze fixation for both verbal-cued and tactile-cued targets, suggesting the use of a vision-dependent exteroceptive body representation. The enhancement of the visual body representations by sensorimotor processes was suggested by the greater pointing accuracy when the vertex was identified by tactile stimulation compared to verbal instruction. Moreover, we found in a control condition that participants were more accurate in indicating the position of their own vertex than the vertex of other people. This result supports the idea that sensorimotor experiences increase the spatial resolution of the exteroceptive body representation. Together, our results suggest that the position of rarely viewed body parts are spatially encoded by an exteroceptive body representation and that non-visual sensorimotor processes are involved in the constructing of this representation.

## Introduction

Our daily experience shows that we can touch any part of our body with our hands. Remarkably, this includes touching regions that we rarely see (e.g., top of the head, back). This capacity provides evidence for the existence of internal body representations and for the access of the arm motor system to these representations.

Much of our knowledge on the control of pointing movements to regions of our body (hereafter named proprioceptive targets) comes from studies in which subjects had to indicate with the index finger different positions on their contralateral arm hidden from view [1–6]. Generally, subjects reach proprioceptive targets fairly accurately (errors < ~2 cm). As a key finding, studies show that the sensory modality used to indicate these targets have an impact on the type of body representation used to encode their position [1,5,7]. More precisely, proprioceptive targets identified by tactile stimuli would be encoded using an interoceptive, somatosensory-based, body representation if the eyes or head remain still between the stimulation and the response. The movements directed toward such tactile-cued somatosensory targets would principally engage a fronto-central cortical network [8]. On the other hand, proprioceptive targets identified by auditory or verbal cues would be encoded in an exteroceptive, visually-based, body representation. In this case, the pointing movements would rely to a greater extent on a parieto-occipital cortical network [5] (see [9] for a review of different body representation taxonomies).

Visual calibration of proprioception is required for building coherent somatosensory body representations [10]. The question then arises as to whether proprioceptive targets which are rarely viewed are also spatially encoded by an interoceptive body representation, even when indicated by tactile stimulation. We addressed this issue by asking adult participants to touch with their index finger different points on their head (notably the topmost position of the head). The spatial position of these points were indicated by the experimenter either verbally or by tactile stimulation. We used an established technique for pinpointing the type of body representation used for the spatial encoding of targets which consisted of assessing the effect of peripheral gaze fixation on the pointing accuracy [2,5,7]. The rationale for using this method is that the encoding of a target position is gaze-dependent when using an exteroceptive, visually-based, representation: in this case, the pointing errors are biased in the opposite direction to the gaze. Inversely, the spatial encoding of a target with an interoceptive body representation would be gaze independent: in this case, the pointing accuracy is not affected by gaze direction.

Clues exist in the literature suggesting that an exteroceptive body representation could be favoured over interoceptive representation for localising rarely viewed proprioceptive targets identified by tactile stimulation. Indeed, studies have shown that coherent visual representations, including of human bodies, can be built using partial visual information [11]. This opens the possibility of building a relatively accurate visual representation using available, yet incomplete visual feedback of the body. Vision of others’ bodies [12] and prior knowledge [13] could contribute to this cognitive construction of body parts that remain out of view.

On the basis of the above mentioned psychophysiological findings, we hypothesized that the spatial positions of body areas which are rarely if ever viewed (e.g., top of the head) are encoded using an exteroceptive body representation, both when they are verbally or tactile-cued. Accordingly, we predicted that the accuracy with which individuals point to these body areas would be biased when gaze direction is deviated from straight-ahead during the pointing movements.

## Methods

### Participants

27 right-handed participants (14 women, mean age: 23.8 ± 2.4 years) volunteered for the experiment. All participants had normal or corrected to normal vision. Informed consent was obtained before the start of the experiment which was carried out in accordance with The Code of Ethics of the World Medical Association (Declaration of Helsinki) for experiments involving humans.

### Experimental set-up

Before running the experiment, the experimenter marked the position of the vertex (i.e. the topmost point of the cranium [14]) on the fabric cap worn by the participant. The vertex is located at the intersection point of the nasion-inion line (fronto-sagittal plane) and of the left and right tragus line (medio-lateral plane) [15]. Then, the participant was seated in semi-darkness with the head vertical and aligned with the trunk. Three light-emitting diodes (LEDs) were positioned eye level ~57 cm in front, at −25 deg (left), 0 deg (central) and +25 deg (right) with respect to the cyclopean eye (Fig. 1). These LEDs served to control gaze direction during the trials. Three small spheres, each 10 cm apart, were fixed on the table, in the participants’ fronto-sagittal plane (the closest at 20 cm from the participants). These spheres served as starting positions for the right index finger. The view of the finger in its initial position helped maintain proprioception calibration during the movement initiation [10].

**Figure 1:**
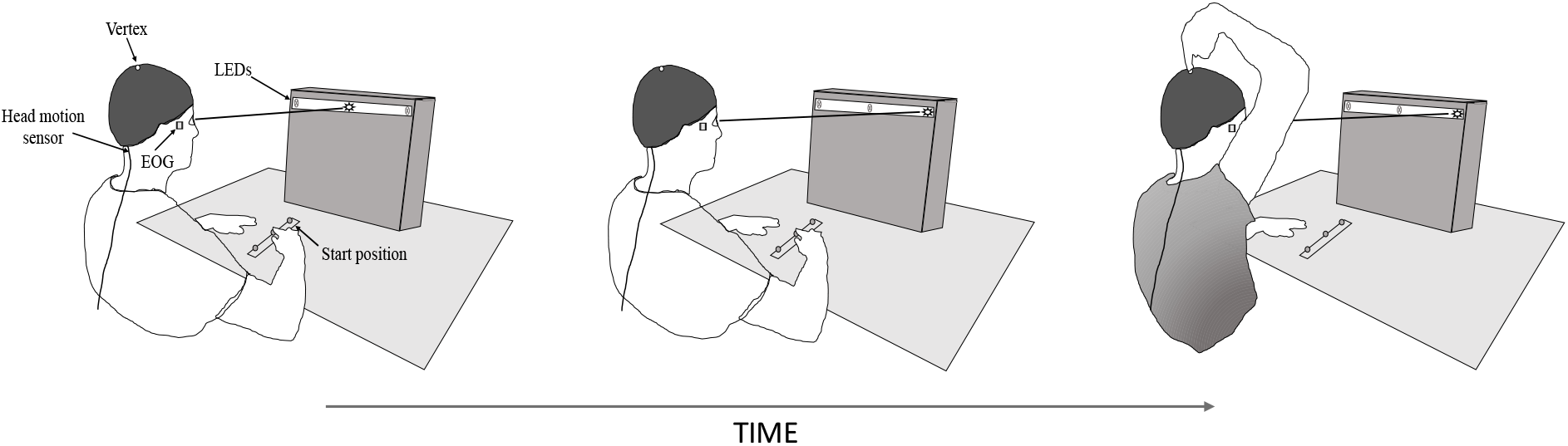
Schematic representation of the experimental set-up and procedure. The figure depicts the time course of a trial involving the middle starting position of the finger and a rightward gaze fixation. EOG: electro-oculography.

### Experimental conditions

The task of the participants was to indicate different points on their head with the tip of the index finger. This task was performed in two conditions (i.e., Verbal, Tactile) which differed according to the cue modality used to indicate these targets.

#### Verbal-cue condition

At the start of each trial, the experimenter verbally indicated one of the 3 finger starting positions (i.e., far, middle or close). After placing the tip of the right index finger on the corresponding home position, the participant sent the verbal message “ready” to the experimenter. Then, the following lighting sequence of the fixation LEDs started. From 0 to 1.5 s: lighting of the central LED; from 2 to 10 s: lighting of either the left, central or right LED. The participant had to fix the lit LED without moving the head. After ensuring that the participant fixated the final lit LED by looking at the electro-oculography (EOG) signal on a computer screen, the experimenter verbally indicated one of the 3 points of the head (hereafter referred to as the targets) that the participants had to reach with the index finger.

The proprioceptive targets were described and named as follows: the topmost point of the head (top), the mid-distance between the eyes and the topmost point of the head (front) and the mid-distance between the rearmost and the topmost points of the head (back). Without moving the eyes or the head, the participants had to touch, as accurately as possible, the target with the tip of their index finger. The participants were instructed that they should not rush their response. Standing behind the participants, the experimenter slipped a digitalizing stylus under the index to record the touched position on the head. After that, the experimenter touched 4 other points of the head with the stylus. The third of which was the actual vertex, while the others were nearby random positions on the head. The participants were told that, after the experiment, they would be informed of the reasons why they were touched at different positions with the stylus. Touching the head at random positions diminished the possibility of the participants obtaining error feedback about their pointing accuracy. Questioned after the experiment, all participants confirmed that they did not realize that the third position touched by the stylus on their head corresponded to the actual target position. Only trials using the vertex as the target were analyzed. The use of the front and back targets, and of the different starting positions minimized the risk of participants implementing stereotyped pointing responses to the vertex. For this reason, the front and back targets were not defined precisely. Note however that the direction of the targets with respect to the initial finger position remained similar in all trials.

#### Tactile-cue condition

This condition was similar to the Verbal-cue condition. Instead of verbally identifying the targets, the experimenter touched the targets using a Semmes-Weinstein monofilament (T = 6.65; force: 2.94 N). To this end, the experimenter held the filament perpendicularly above the target before descending it until the filament bowed upon contact with the head. This contact was held steady for ~1s. After the tactile stimulation, the participants touched with the tip of the index finger where they felt they had been touched on their head. The use of the Semmes-Weinstein monofilament ensured that the sensory stimulation evoked by the touches was similar across conditions (i.e., gaze direction and cue condition) and across participants. Although the somatosensory threshold over the vertex is higher compared to over other regions of the scalp and body [16], all participants clearly perceived the touches. Note that this condition was named “Tactile-cue” despite that the hair follicles are innervated with Merkel disks and lanceolate nerve endings [17] Touching the head cap with the filament therefore created a somatosensory amalgam providing spatial information of the stimulation.

For both the Cue (Verbal, Tactile) and Gaze (Left, Central, Right) conditions, 12 trials used the vertex target, 2 trials used the front target and 2 trials used the back target. The order of presentation of the targets and of the fixation LED was pseudorandom. 14 participants started the experiment with the Tactile-cue condition. Prior to each experimental condition, participants performed a series of 7 familiarization trials comprising all combinations of ocular fixation with the front and back targets (i.e. 6 trials) and 1 trial using the top target and the central fixation.

#### Others (control condition)

We designed a control condition (referred to as Others) to assess the accuracy with which one can localize the vertex of other people. The identification of this position can be thought of as being essentially based on the visual image of another person’s head. The participants located with the tip of their index finger the vertex of 6 adult volunteers (aged between 21 and 56 years) wearing a black fabric cap. During the test, the volunteers were seated, eyes closed, in a lighted room with the head vertical and aligned with the trunk. The experimenter positioned the tip of the digitizing stylus at the point of the head indicated by the participants. The participants were then asked to move around the seated volunteer to evaluate whether the stylus truly indicated the vertex. They were allowed to correct the position of the stylus, before the experimenter marked the position touched by the stylus. Then, in the absence of the participants, the actual vertex position of the volunteers was also marked on the cap before this and the actual vertex positions were recorded with the stylus. This control condition was performed either before or after both the Verbal-cue and Tactile-cue conditions.

### Data recordings

The endpoint position of the index finger on the head was recorded using a digitizing stylus (120 Hz, Polhemus Fastrak, Vermont, USA). Head position was tracked using an electromagnetic sensor (120 Hz, Polhemus Fastrak, Vermont, USA) embedded in the cap at the back of the head. The position of the eyes was recorded at 250Hz by EOG (Coulbourn Instruments, Lehigh Valley, PA). The EOG recordings were displayed on the computer screen. They were used to verify the participant’s ocular behavior during the experiment. The few trials in which the participants failed to maintain their ocular deviation before or during their pointing movement were deleted and repeated.

#### Data analyzes

The perception of the vertex position was assessed by computing the radial error, which was defined as the difference between the participant’s finger end position and the actual vertex position (i.e., the 2D euclidean distance). Previous studies have shown that gaze direction had a greater impact on pointing movements in the medio-lateral direction than in the antero-posterior direction when subjects have their head aligned with the trunk as in the present study [2,5,7]. The effect of Gaze and Cue-target conditions on the perception of the vertex was then further analyzed by computing the lateral (X coordinates) and longitudinal (Y coordinates) errors. These errors were respectively defined as the signed distance between the perceived and the actual vertex positions in both X and Y coordinates. Positive lateral errors indicated that the participants perceived the vertex to the right of the actual vertex. Positive longitudinal errors indicated that the vertex was perceived in front of the actual vertex.

The variability (i.e., standard deviation of the mean) in estimating the vertex position (i.e., radial, lateral and longitudinal errors) was computed for each Gaze and Cue-target conditions. This variable provides an estimate of the reliability of the representation used to locate the vertex.

The continuous recording of the head position indicated that the participants succeeded in minimizing head movements between the recordings of their touched position and of the actual vertex position. On average (all Cue-target and Gaze conditions), the participants moved their head by 0.124 ± 0.274 cm in the transverse plane between the two recordings. To cancel out the effect of these small head movements on the assessment of the participants’ performance, for each trial, we subtracted X (medio-lateral) and Y (antero-posterior) head displacements measured between the two recordings, from the vertex X and Y data prior to the error calculation.

Statistical analyses were carried out with the Statistica software (Statsoft, inc). The effect of Gaze and of Cue-target conditions was tested using 3 Gaze (left, center, right) x 2 Cue-target (Verbal, Tactile) repeated measures analyses of variance (ANOVAs). We also compared the accuracy with which participants perceived the position of their own vertex with the accuracy which they perceived the position others’ vertex. To this end, the mean errors (radial, lateral, longitudinal) measured while the participants gazed at the central LED in both the Verbal-cue and Tactile-cue conditions were compared with those measured in the Others condition. The data were submitted to separate one-way ANOVAs. The alpha level was set at 0.05 for all statistical contrasts. Significant effects were further analyzed using Newman-Keuls post-hoc tests.

## Results

### Radial error

As a first salient finding, the amplitude of the radial error significantly differed according to the type of cue indicating the vertex (F_1,26_ = 28.29; p < 0.001, Fig. 2A). The radial error was smaller in the Tactile-cue (1.04 ± 0.44 cm) than in the Verbal-cue (1.38 ± 0.57 cm) conditions. The ANOVA did not reveal significant effect of Gaze (F_2,26_ = 0.13; p = 0.88) or significant interaction Cue-Target x Gaze (F_2,26_ = 0.42; p = 0.66).

**Figure 2:**
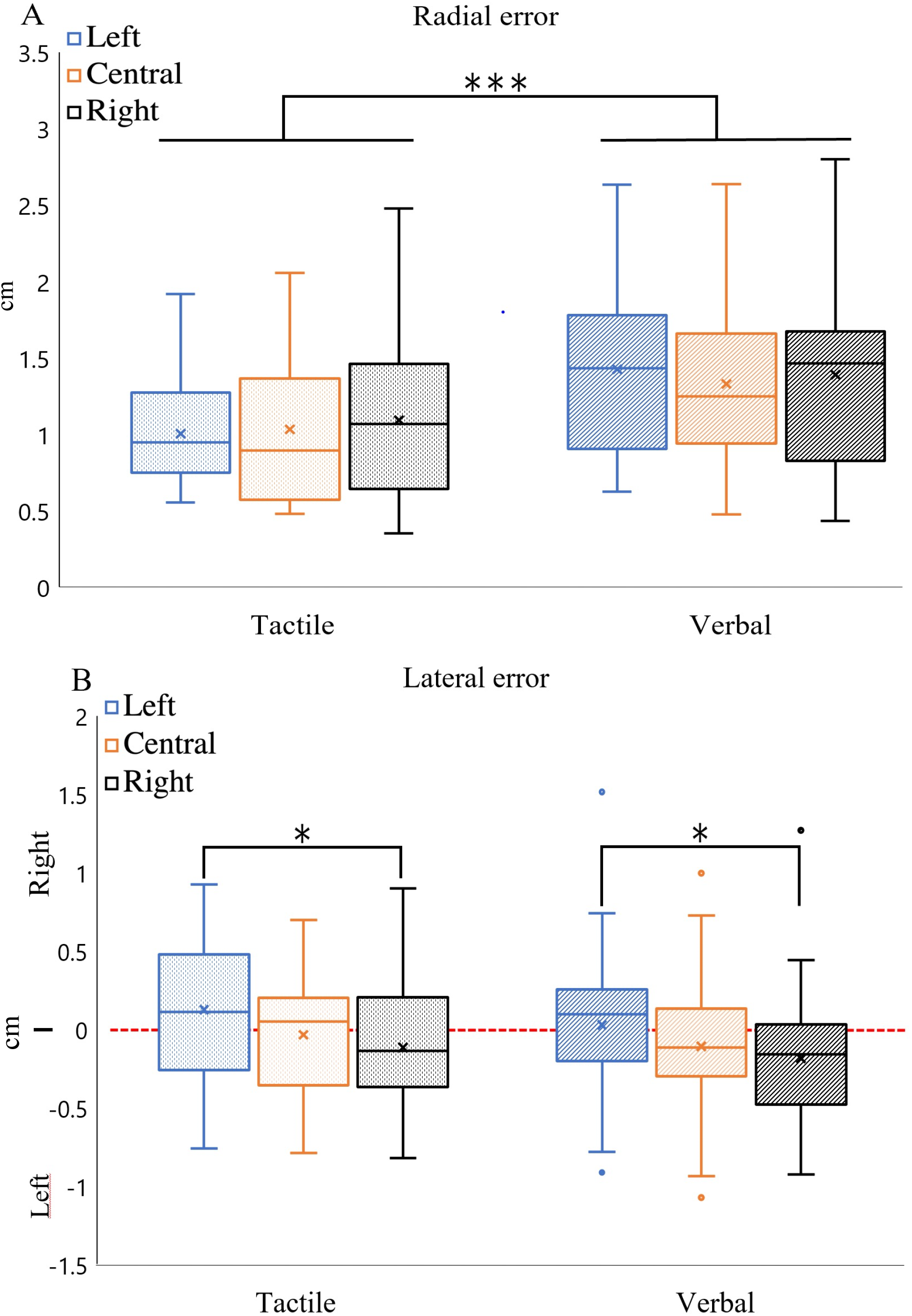
Boxplots of radial [A] lateral [B] errors measured in the different Cue-target and Gaze conditions.*p = 0.02; ***p < 0.001.

The variability of the radial error was significantly different between the two Cue-target conditions (F_1,26_ = 27.94; p < 0.001). This variability was smaller when the participants indicated their vertex position in the Tactile-cue (0.70 ± 0.28 cm) than in the Verbal-cue (0.88 ± 0.30 cm) conditions. The analyses of the radial error variability yielded no significant effect of Gaze (F_2,26_ = 0.24; p = 0.79), and no significant interaction Cue-target x Gaze (F_2,26_ = 0.09; p = 0.91).

### Lateral error

The participants perceived their vertex slightly to the left of its actual position with right gaze fixation (0.11 ± 0.48 cm) and slightly to the right with left gaze fixation (−0.15 ± 0.48 cm). This bias was confirmed by the ANOVA which revealed a significant effect of Gaze on the lateral error (F_2,26_ = 4.20; p = 0.02. Fig. 2B) and by the post-hoc comparison which showed a significant difference between left and right gaze fixations (p = 0.02). The perceived vertex position in the gaze-centered condition did not significantly differ from that measured in gaze-deviated conditions (both p > 0.05). Importantly, the perceived lateral position of the vertex did not significantly differ between the Verbal-cued and the Tactile-cued target conditions (F_1,26_ = 3.42; p = 0.07) and the interaction Gaze x Cue-target was not significant (F_2,26_ = 0.03; p = 0.97).

The variability of the lateral error was significantly different between the two Target-cue conditions (F_2,26_ = 15.86; p < 0.001). Indeed, this variability was smaller in the Verbal-cue condition (0.46 ± 0.14 cm) compared to the Tactile-cue condition (0.56 ± 0.22 cm). In contrast, there was no significant effect of Gaze (F_2,26_ = 0.19; p = 0.82) and no significant interaction Gaze x Cue-target (F_2,26_ = 0.08; p = 0.93).

### Longitudinal error

The longitudinal error was not significantly different between the Verbal-cue and Tactile-cue conditions (F_2,26_ < 0.001; p = 0.98) or between the different gaze fixations (F_2,26_ = 0.79; p = 0.46). The interaction Gaze x Cue-target was also not significant (F_2,26_ = 0.49; p = 0.61).

The variability of longitudinal error was significantly different between the two Target-cue conditions (F_1,26_ = 9.30; p = 0.003). Indeed, this variability was smaller in the Tactile-cue condition (1.05 ± 0.37 cm) compared to the Verbal-cue condition (1.22 ± 0.46 cm). In contrast, there was no significant effect of Gaze (F_2,26_ = 0.01; p = 0.99) and no significant effect of interaction Gaze x Cue-target (F_2,26_ = 0.70; p = 0.50).

### Perceived self versus others vertex position

#### Radial error

The participants’ perception of the position of another person’s’ vertex was compared with the perception of their own vertex position in the Tactile and Verbal conditions. For this comparison, we used the errors (i.e., radial, lateral and longitudinal) measured in both Cue-target conditions while the participants were looking straight-ahead. The one-way ANOVA (Others, Tactile, Verbal) yielded a significant effect of Condition on the radial errors (F_2,26_ = 15.27; p < 0.001, Fig. 3). Post-hoc comparison revealed that the radial error significantly differed between each condition. The radial errors were greatest in the Others condition (1.83 ± 0.71 cm) and smallest in the Tactile-cue conditions (1.03 ± 0.46 cm). As it can be seen in Fig. 3, two participants were identified as outliers in the boxplot of the radial errors in the Others condition. To ensure that the larger radial error in the Others condition was not due to these participants, we performed another ANOVA without them. The significant effect of condition was preserved (F_2,24_ = 13.09; p < 0.001) as well as all pairwise comparisons (all p < 0.05).

**Figure 3:**
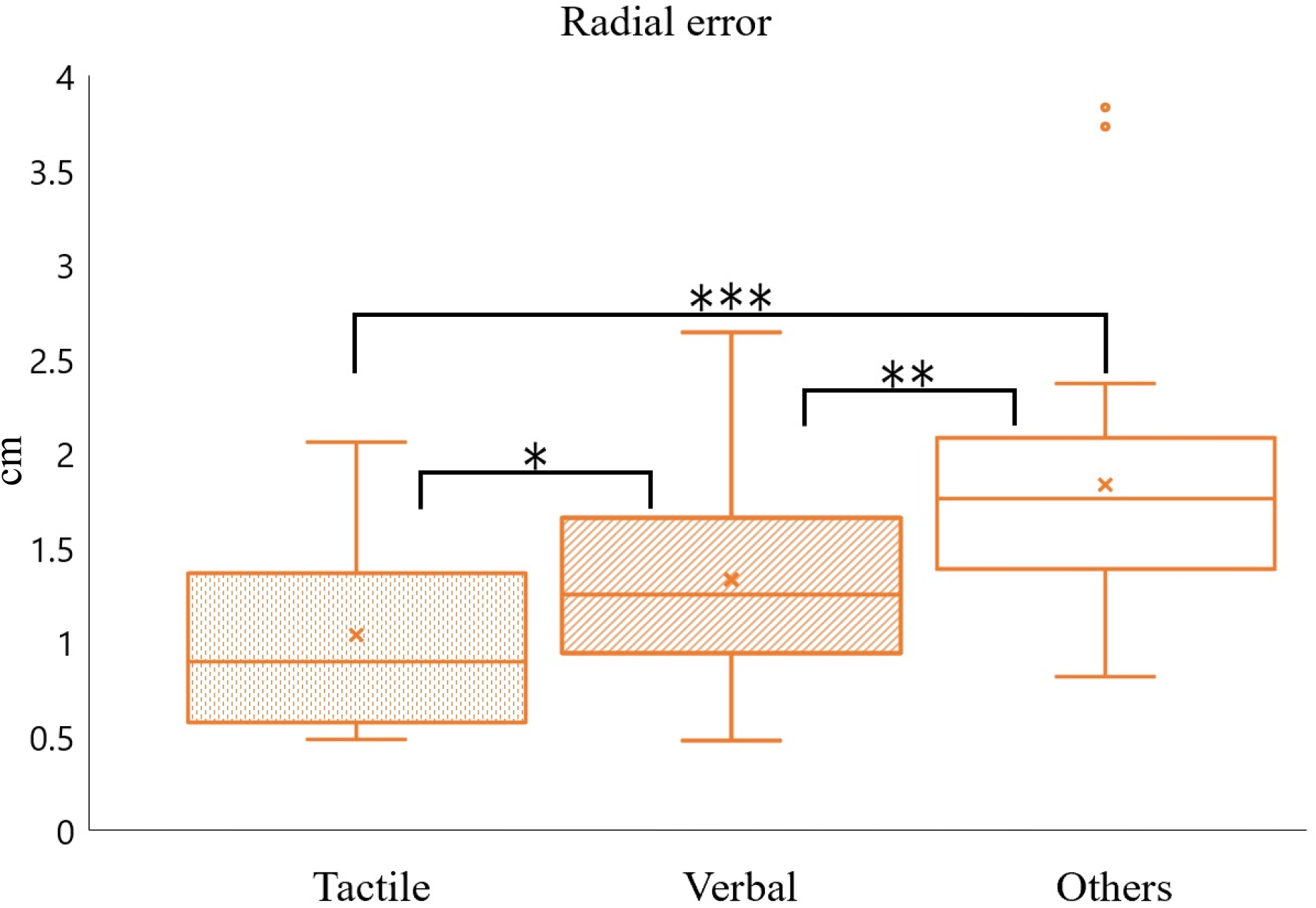
Boxplot of radial errors in the Tactile-cued condition and in the Verbal-cued and Others conditions (both with centered-gaze fixation).*p = 0.05; **p = 0.005; ***p < 0.001

### Lateral and longitudinal errors

The ANOVA revealed a significant effect of Condition on the lateral error (F_2,26_ = 4.05; p = 0.02). The Post-hoc analyses showed that the lateral error in the Others condition (−0.29 ± 0.45 cm) was significantly greater than in the Tactile-cue condition (0.01 ± 0.35 cm, p = 0.02), but not significantly different to the Verbal-cue condition (−0.13 ± 0.48 cm, p = 0.09).

The longitudinal error did not significant differ (F_2,26_ = 1.50; p = 0.23) between Tactile-cue (−0.02 ± 1.06 cm), Verbal-cue (0.10 ± 1.35 cm), and Others conditions (0.51 ± 1.16 cm).

## Discussion

Previous studies have shown that the sensory modality used to identify the position of proprioceptive targets influences the type of the body representation employed for reaching them with the finger [1,5]. More specifically, identifying the target with an auditory and tactile cue respectively prompts the use of an exteroceptive (visually-based) and interoceptive (somatosensory-based) body representations when the eyes and head remain stationary. This scenario appears to be less straightforward when the proprioceptive target consists in a rarely, if ever, viewed area of the body, as in the present study. Indeed, when pointing to the tactile-cued vertex, our participants presented gaze-dependent errors. This effect of gaze direction on pointing accuracy is consistent with the use of an exteroceptive body representation [2,5,7]. One interpretation for this novel finding could be that visual body representations are used to localize body parts that cannot be identified through visual-proprioceptive integration. We shall focus much of our discussion on the bases of this interpretation.

Whilst the proprioceptive targets used in previous studies were generally located on the contralateral arm [1–6], our participants had to indicate the position of their vertex with the tip of their finger. The peculiarity of this body location is that it is rarely, if ever, viewed on ourselves. Our results therefore lead to the paradox that the spatial position of a tactile stimulation on a part of the body that is regularly viewed would be encoded through a somatosensory-based representation [1,3,5] while the position of a touch on a rarely-viewed body part would be encoded through a visually-based representation (present results). This apparent paradox could be resolved by considering that the intrinsic and extrinsic body representations are co-constructed [13] and that the weight given to each representation is context-dependent.

An accurate somatosensory mapping of the body space requires cross sensory calibration [18]. For sighted persons, this calibration is principally achieved through vision [10,13]. The possibility to regularly see our upper limbs and therefore refresh their somatosensory mapping through vision could enhance the reliability of the intrinsic body representation for encoding the spatial position of tactile stimuli on the arm. The reliability of the intrinsic body representation for encoding the spatial position of hidden body parts such as the vertex, appears hampered by the impossibility to co-register their positions with somatosensory and visual inputs. The lack of visual calibration could also have an exacerbated detrimental effect for localizing body areas with little density of somatosensory receptors, as is the case for the top of the head [16].

In this context, an extrinsic body representation appears most suitable for coding the vertex position. Seeing ourselves from a first or a third (e.g., through a mirror) perspective, and seeing someone else’s body [12] would be fundamental to constructing what has been referred to as the long-term visual body representation (for a review, see [13]). This body representation could also benefit from the capacity of the brain to construct coherent consolidated representations, or to re-actualise them, on the basis of partial visual information [11].

Furthermore, sensorimotor-derived processes could also be involved in the construction of visual body representations [19]. For instance, sensorimotor experience gained during a lifetime could have a key role in enhancing the spatial resolution of the visual representation of the vertex. Putting on a sweatshirt or a hat, or washing your hair are examples of sensorimotor activities that might improve the visual representation of this unseen body area. This idea is supported by the fact that our participants showed smaller radial and lateral errors when indicating the position of their own vertex, than the vertex of other people. It seems reasonable to assume that the task of localizing the vertex on another person’s head is essentially based on visual representations despite that our own body representations, including those somatosensory-based, may intervene in identifying others’ body parts [20]. Our results therefore further stress the importance of non-visual sensorimotor processes for constructing body representations, including those of visual origin. This is in line with the proposal that intrinsic and extrinsic body representations are co-constructed [13].

The enhancement of visual body representations by sensorimotor processes is also suggested by the greater pointing accuracy showed by the participants when their vertex was identified by tactile stimulation compared to verbal instruction (i.e. smaller radial error associated with smaller variability). Previous studies have reported improved position sense when applying tactile stimulation on the proprioceptive targets [4,6]. Our results indicate that this finer spatial resolution of the somatosensory mapping (which could be short-lived) can benefit visually-based extrinsic body representations. The interdependence between somatosensory-based and visually-based body representations is also revealed in the so-called rubber hand illusion [21]. This illusion arises from the simultaneous brushing of the hand of the subject, which is hidden from view, and of a facsimile of a human hand viewed in front of the subject. After a few minutes of exposure to this somato-visual context, the subjects perceive the fake hand as being the real hand, consistent with a close link between visual and somatosensory body representations.

## Acknowledgments

The authors thank Fanny Goetz for her help at various stages of this research project.

## Declarations of interest

none

## Notes

### Competing Interest Statement

The authors have declared no competing interest.

## References

[1] A. Sirigu, J. Grafman, K. Bressler, T. Sunderland, Multiple representations contribute to body knowledge processing: evidence from a case of autotopagnosia, Brain. 114 (1991) 629–642. https://doi.org/10.1093/brain/114.1.629.

[2] A. Blangero, Y. Rossetti, J. Honoré, L. Pisella, Influence of gaze direction on pointing to unseen proprioceptive targets, Adv. Cognit. Psychol. 1 (2005) 9 16. https://doi.org/10.2478/v10053-008-0039-7.

[3] F.R. Sarlegna, R.L. Sainburg, The effect of target modality on visual and proprioceptive contributions to the control of movement distance, Exp. Brain Res. 176 (2007) 267–280. https://doi.org/10.1007/s00221-006-0613-5.

[4] L. Mikula, S. Sahnoun, L. Pisella, G. Blohm, A.Z. Khan, Vibrotactile information improves proprioceptive reaching target localization, Plos One. 13 (2018) e0199627. https://doi.org/10.1371/journal.pone.0199627.

[5] G.A. Manson, L. Tremblay, N. Lebar, J. de Grosbois, L. Mouchnino, J. Blouin, Auditory cues for somatosensory targets invoke visuomotor transformations: Behavioral and electrophysiological evidence, Plos One. 14 (2019) e0215518. https://doi.org/10.1371/journal.pone.0215518.

[6] A. Goettker, K. Fiehler, D. Voudouris, Somatosensory target information is used for reaching but not for saccadic eye movements, J. Neurophysiol. 124 (2020) 1092–1102. https://doi.org/10.1152/jn.00258.2020.

[7] S. Mueller, K. Fiehler, Effector movement triggers gaze-dependent spatial coding of tactile and proprioceptive-tactile reach targets, Neuropsychologia. 62 (2014) 184–193. https://doi.org/10.1016/j.neuropsychologia.2014.07.025.

[8] P.-M. Bernier, B. Burle, T. Hasbroucq, J. Blouin, Spatio-temporal dynamics of reach-related neural activity for visual and somatosensory targets, Neuroimage. 47 (2009) 1767–1777. https://doi.org/10.1016/j.neuroimage.2009.05.028.

[9] F. de Vignemont, Body schema and body image—Pros and cons, Neuropsychologia. 48 (2010) 669 680. https://doi.org/10.1016/j.neuropsychologia.2009.09.022.

[10] V. Harrar, L.R. Harris, Eye position affects the perceived location of touch, Exp. Brain Res. 198 (2009) 403–410. https://doi.org/10.1007/s00221-009-1884-4.

[11] C.T. Fuentes, M.R. Longo, P. Haggard, Body image distortions in healthy adults, Acta Psychol. 144 (2013) 344–351. https://doi.org/10.1016/j.actpsy.2013.06.012.

[12] S. Gallagher, How the Body Shapes the Mind, Oxford University Press, 2005. https://doi.org/10.1093/0199271941.001.0001.

[13] V. Pitron, F. de Vignemont, Beyond differences between the body schema and the body image: insights from body hallucinations, Conscious. Cogn. 53 (2017) 115–121. https://doi.org/10.1016/j.concog.2017.06.006.

[14] D. Kempe, M. Salek, C. Moore, Frugal and truthful auctions for vertex covers, Flows and Cuts, in: 2010 IEEE 51st Annual Symposium on Foundations of Computer Science, IEEE, Las Vegas, NV, USA, 2010: pp. 745–754. https://doi.org/10.1109/FOCS.2010.76.

[15] R.W. Homan, J. Herman, P. Purdy, Cerebral location of international 10–20 system electrode placement, Electroencephalogr. Clin. Neurophysiol. 66 (1987) 376–382. https://doi.org/10.1016/0013-4694(87)90206-9.

[16] A. Mehrabyan, S. Guest, G. Essick, F. McGlone, Tactile and thermal detection thresholds of the scalp skin, Somatosens. Mot. Res. 28 (2011) 31–47. https://doi.org/10.3109/08990220.2011.602764.

[17] K. Taira, Y. Narisawa, J. Nakafusa, N. Misago, T. Tanaka, Spatial relationship between Merkel cells and Langerhans cells in human hair follicles, J. Dermatol. Science. 30 (2002) 195–204. https://doi.org/10.1016/S0923-1811(02)00104-4.

[18] R.J. van Beers, D.M. Wolpert, P. Haggard, When feeling is more important than seeing in sensorimotor adaptation, Curr. Biol. 12 (2002) 834–837. https://doi.org/10.1016/S0960-9822(02)00836-9.

[19] F. Mancini, M.R. Longo, G.D. Iannetti, P. Haggard, A supramodal representation of the body surface, Neuropsychologia. 49 (2011) 1194–1201. https://doi.org/10.1016/j.neuropsychologia.2010.12.040.

[20] H. Ishida, K. Nakajima, M. Inase, A. Murata, Shared mapping of own and others’ bodies in visuotactile bimodal area of monkey parietal cortex, J. Cogn. Neurosci. 22 (2010) 83–96. https://doi.org/10.1162/jocn.2009.21185.

[21] H.H. Ehrsson, C. Spence, R.E. Passingham, That’s my hand! Activity in premotor cortex reflects feeling of ownership of a limb, Science. 305 (2004) 875–877. https://doi.org/10.1126/science.1097011.

